# Seqping: Gene Prediction Pipeline for Plant Genomes using Self-Trained Gene Models and Transcriptomic Data

**DOI:** 10.1101/038018

**Authors:** Kuang-Lim Chan, Rozana Rosli, Tatiana Tatarinova, Michael Hogan, Mohd Firdaus-Raih, Eng-Ti Leslie Low

**Affiliations:** Malaysian Palm Oil Board, 6, Persiaran Institusi, Bandar Baru Bangi, 43000 Kajang, Selangor, Malaysia; Keck School of Medicine, University of Southern California, Los Angeles, CA, USA; Orion Genomics, 4041 Forest Park Avenue, St. Louis, MO 63108, USA; School of Biosciences and Biotechnology, Faculty of Science and Technology, and Institute of Systems Biology, Universiti Kebangsaan Malaysia, 43600 Bangi, Selangor, Malaysia

## Abstract

**Summary:** Although various software are available for gene prediction, none of the currently available gene-finders have a universal Hidden Markov Models (HMM) that can perform gene prediction for all organisms equally well in an automatic fashion. Here, we report an automated pipeline that performs gene prediction using self-trained HMM models and transcriptomic data. The program processes the genome and transcriptome sequences of a target species through GlimmerHMM, SNAP, and AUGUSTUS training pipeline that ends with the program MAKER2 combining the predictions from the three models in association with the transcriptomic evidence. The pipeline generates species-specific HMMs and is able to predict genes that are not biased to other model organisms. Our evaluation of the program revealed that it performed better than the use of the closest related HMM from a standalone program.

**Availability and Implementation:** Distributed under the GNU license with free download at http://sourceforge.net/projects/seqping and http://genomsawit.mpob.gov.my.

**Supplementary information:** Supplementary data are available at Bioinformatics online.

## 1 Introduction

Next-generation sequencing technologies are producing large volumes of sequence data for many large-scale genome projects. As the volume of sequence data increases exponentially, the task of efficient genome annotation becomes especially critical. Biological interpretation and annotation are time consuming and labour intensive. There is a need for accurate and fast tools to analyze these sequences, especially to identify genes and determine their functions. Algorithms and pipelines, such as MAKER2 (Holt and Yandell, 2011), Fgenesh++ (Solovyev *et al*., 2006), GeneMark.hmm (Lomsadze *et al*., 2005), GlimmerHMM (Majoros *et al*., 2004), AUGUSTUS (Stanke *et al*., 2006), and SNAP (Korf, 2004) have been developed with the intention to address this problem. Never theless, many of the currently available gene finders base their predictions on known HMM models and species-specific parameters, leading to biases in the gene prediction (Sleator, 2010). This reduces the accuracy of the gene models predicted. Thus, predicting the protein-coding genes remains a complex and significant challenge.

Here we present a gene prediction pipeline that generates species-specific HMMs for use on any newly sequenced plant genome. Our program, Seqping, combines existing gene finders with self-trained HMMs constructed from a training set of the same species. This pipeline automates and streamlines the gene prediction process by preparing the training data, building HMMs, and performing gene prediction.

## 2 Methods

Seqping is a command-line executed shell program to automate the gene prediction process by executing a sequence of commands and other shell and Perl scripts (Supplementary Figure 1). The Seqping pipeline is divided into six stages: (i) preparation of working directories; (ii) processing of transcriptome sequences; (iii) GlimmerHMM training and prediction; (iv) AUGUSTUS training; (v) SNAP training, and (vi) MAKER2 prediction. Seqping is built with options to utilize multiple processors and distribute job submissions to cluster nodes.

The Seqping program requires the user to submit transcriptome and genome sequences of the target species, as well as a Reference Proteins (RF) set in FASTA format. The RF dataset is composed of selected protein sequences of full-length coding sequences (CDS) from RefSeq (Pruitt *et al*., 2012) and UniProtKB (The UniProt Consortium, 2013). For the experiments described here, only protein coding sequences for species under the phylum of magnoliophyta or flowering plant were used. The RF dataset was filtered to exclude hypothetical proteins, ribosomal proteins, tRNAs, mitochondrial and chloroplast proteins. The TIGR Plant Repeat (Ouyang *et al*., 2004) and RepBase (Jurka *et al*., 2005) sequences were combined to a single FASTA file for additional TBLASTX (Altschul *et al*., 1990) filtering. HMM profiles from the Gypsy Database (Llorens *et al*., 2011) were used for HMMER (Johnson *et al*., 2010) hmmsearch filtering. The final tool in the pipeline, MAKER2, combines all the predictions and provides GFF3 formatted outputs, as well as the predicted genes and proteins in FASTA format. A comprehensive log file is generated concurrently while executing the program.

### 2.1 Training Set Preparation

Seqping first extracts the open reading frames (ORFs), sized between 500 and 5000 nucleotides, from the transcriptome set using *getorf* from the EMBOSS package (Rice *et al*., 2000). Next, the ORFs with RF support (BLASTX; E value of 1e-10 or lower) are stringently clustered using BLASTClust and CD-HIT-EST (Fu *et al*., 2012). After repeats filtering, the final sequences were used as the training set to develop species-specific HMMs for gene prediction.

### 2.2 HMM Training

In this step, the program aligns the training set against the genome using Splign and Compart (Kapustin *et al*., 2008). The aligned training set with their respective genome sequences were used to train GlimmerHMM. In Seqping, a Perl script converts the Splign output into an exon file, and then runs trainGlimmerHMM to produce a HMM model. Gene prediction by GlimmerHMM is executed using the species-specific HMM model, followed by repeats filtering.

The training set for AUGUSTUS is translated into protein sequences using EMBOSS’s *transeq.* A new HMM is produced using an AUGUSTUS specific training script that can be found in the AUGUSTUS package.

In order to build a HMM for SNAP, Seqping runs a basic MAKER2 prediction using nucleic acids and protein sequences from the training set. The SNAP HMM model is finally produced by *fathom* and *hmm-assembler* scripts from the SNAP package.

### 2.3 Final Prediction

MAKER2 takes as input the GFF3 file of GlimmerHMM, AUGUSTUS HMM, and SNAP HMM, in addition with the transcriptome data, for the final gene prediction step. To make the process run in parallel, the genome sequences are split into multiple files according to the number of CPU defined by user.

## 3 Results

By training species-specific HMMs, Seqping provides an effective, organism independent, gene prediction tool for non-model plant species. Expectedly, the performance is influenced by the quality of transcriptome and genome sequences of the target species.

### 3.1 *Oryza sativa* Gene Prediction

The pipeline was tested by simulating the gene prediction process of the rice *(Oryza sativa ssp. japónica)* genome. The fourteen genomic sequences used are rice genome pseudomolecules from the MSU Rice Genome Annotation Project release 7 (Kawahara *et al.,* 2013). The transcriptome set used were the assembled transcripts from three RNA-Seq projects in NCBI BioProject: PRJNA79825, PRJDA67119, and PRJNA80103. A total of 175,251 assembled transcripts were used as input for the pipeline. The contigs N50 and mean length are 1693 and 956 respectively.

The transcripts were treated as described in the Methods section, resulted in 11,729 putative full-length ORFs that were used for HMM training. The MAKER program was then invoked to combine the predictions from the three modelers with the transcrip-tomic data as evidence. The Seqping pipeline identified 24,009 highly confident genes.

It took approximately 100 hours to execute the gene prediction pipeline on a Linux cluster with 9 nodes (8 CPUs per node). The predicted genes, when measured by comparing them to the MSU annotation using ParsEval (Standage and Brendel, 2012), yielded 87.70% shared gene loci. A total of 24,229 complete genes of *O. sativa ssp. japonica* from RefSeq were used as reference to calculate sensitivity (Sn) and specificity (Sp) as described by Burset and Guigo (1996) using GenomeTools *gt-eval* (Gremme *et al*., 2013). Performance of Seqping was compared to Fgenesh pipeline and three HMM-based programs MAKER, GlimmerHMM and AUGUSTUS (Table 1), and we conclude that the Seqping pipeline predictions are more accurate than gene predictions using the other three approaches with the default or available HMMs.

**Table 1.**
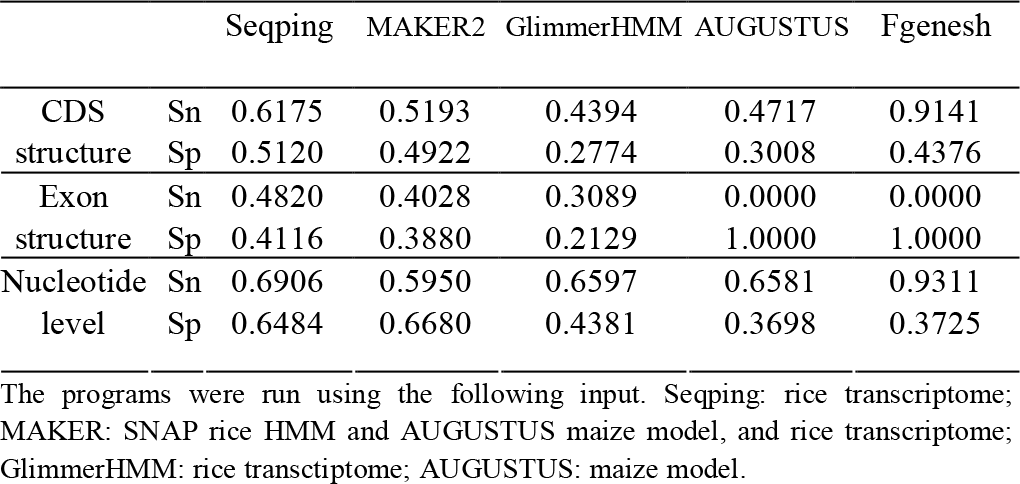
Comparison of gene prediction approaches on *O. sativa*

|  |  | Seqping | MAKER2 | GlimmerHMM | AUGUSTUS | Fgenesh |
| --- | --- | --- | --- | --- | --- | --- |
| CDS | Sn | 0.6175 | 0.5193 | 0.4394 | 0.4717 | 0.9141 |
| structure | Sp | 0.5120 | 0.4922 | 0.2774 | 0.3008 | 0.4376 |
| Exon | Sn | 0.4820 | 0.4028 | 0.3089 | 0.0000 | 0.0000 |
| structure | Sp | 0.4116 | 0.3880 | 0.2129 | 1.0000 | 1.0000 |
| Nucleotide | Sn | 0.6906 | 0.5950 | 0.6597 | 0.6581 | 0.9311 |
| level | Sp | 0.6484 | 0.6680 | 0.4381 | 0.3698 | 0.3725 |

The programs were run using the following input. Seqping: rice transcriptome; MAKER: SNAP rice HMM and AUGUSTUS maize model, and rice transcriptome; GlimmerHMM: rice transcriptome; AUGUSTUS: maize model.

### 3.2 *Arabidopsis thaliana* Gene Prediction

Annotation from *Arabidopsis thaliana* TAIR10 (Lamesch *et al.,* 2011) was also used to compare the performance of Seqping with other programs (Table 2). The genome and transcriptome used were downloaded from TAIR10.

**Table 2.** Comparison of gene prediction approaches on *A. thaliana*

|  |  | Seqping | MAKER2 | GlimmerHMM | AUGUSTUS |
| --- | --- | --- | --- | --- | --- |
| CDS | Sn | 0.2749 | 0.0738 | 0.2804 | 0.3075 |
| structure | Sp | 0.8867 | 0.8877 | 0.7515 | 0.7527 |
| Exon | Sn | 0.4596 | 0.1207 | 0.0000 | 0.4155 |
| structure | Sp | 0.7313 | 0.7457 | 1.0000 | 0.5373 |
| Nucleotide | Sn | 0.7929 | 0.1932 | 0.9350 | 0.9634 |
| level | Sp | 0.9748 | 0.9750 | 0.8150 | 0.7974 |

The programs were run using the following input. Seqping: *A. thaliana* transcriptome; MAKER: SNAP *A. thaliana* HMM and AUGUSTUS *A. thaliana* model, and *A. thaliana* transcriptome; GlimmerHMM: *A. thaliana* transcriptome; AUGUSTUS: *A. thaliana* model.

## Acknowledgements

We thank the Director-General of Malaysian Palm Oil Board (MPOB) for publication authorization and all members of the MPOB Bioinformatics Unit for helpful discussions.

## Funding

MPOB Oil Palm Genome Programme (R009611000).

